# ATLAS: A rationally designed anterograde transsynaptic tracer

**DOI:** 10.1101/2023.09.12.557425

**Authors:** Jacqueline F. Rivera, Weiguang Weng, Haoyang Huang, Sadhna Rao, Bruce E. Herring, Don B. Arnold

## Abstract

Neural circuits, which constitute the substrate for brain processing, can be traced in the retrograde direction, from postsynaptic to presynaptic cells, using methods based on introducing modified rabies virus into genetically marked cell types. These methods have revolutionized the field of neuroscience. However, similarly reliable, transsynaptic, and non-toxic methods to trace circuits in the anterograde direction are not available. Here, we describe such a method based on an antibody-like protein selected against the extracellular N-terminus of the AMPA receptor subunit GluA1 (AMPA.FingR). ATLAS (<u>A</u>nterograde <u>T</u>ranssynaptic <u>L</u>abel based on <u>A</u>ntibody-like <u>S</u>ensors) is engineered to release the AMPA.FingR and its payload, which can include Cre recombinase, from presynaptic sites into the synaptic cleft, after which it binds to GluA1, enters postsynaptic cells through endocytosis and subsequently carries its payload to the nucleus. Testing in vivo and in dissociated cultures shows that ATLAS mediates monosynaptic tracing from genetically determined cells that is strictly anterograde, synaptic, and non-toxic. Moreover, ATLAS shows activity dependence, which may make tracing active circuits that underlie specific behaviors possible.

## MAIN TEXT

Neural circuits carry information through the nervous system, from presynaptic to postsynaptic cells, in a direction referred to as anterograde. Over the years, molecular methods to trace circuits have been developed, which include a widely adopted method based on a glycoprotein (G)-deleted rabies virus that can trace circuits in the retrograde direction and has revolutionized the field of neuroscience [1, 2]. However, developing similar methods to trace circuits anterogradely has proven more challenging [3]. Several methods for anterograde tracing have been introduced; however, none can efficiently trace monosynaptically from genetically determined cells, only in the anterograde direction and without toxicity [3]. In one method, high titer Adeno-Associated Virus serotype one engineered to carry Cre (AAV1-Cre) was used to infect source neurons in a particular brain region [4, 5]. Infecting the brain region containing postsynaptic neurons with a virus carrying a floxed reporter (AAV8-DIO-GeneX) results in the expression of GeneX in cells in that region. Similarly, injections of AAV1-Cre into specific brain regions in transgenic mice expressing floxed reporters lead to the expression of reporters in neurons in downstream regions. This system has low toxicity, is monosynaptic, and can mark postsynaptic cells throughout the brain when used with transgenic animals. However, because of its dependence on the vagaries of the virus, it suffers from several limitations. For example, it labels retrogradely connected neurons in addition to its anterograde targets [4], and it has not been demonstrated to be purely transsynaptic. Moreover, it has not been shown to work with genetically specified starter neurons, which is essential for describing cell-type specific projections in the nervous system.

More recently, a transneuronal tracing system was developed based on an inactivated variant of the yellow fever virus (YFV) used in vaccinations [6]. The original YFV variant has three limitations: It is toxic, moves retrogradely, and can jump multiple synapses. However, eliminating genes encoding the structural proteins C, M, and E, and NS1, which is necessary for replication, can mitigate these problems. The lack of CME can prevent the virus from transmitting beyond the starter cell, and further controlling the amount of NS1 present with a Dox-dependent system can reduce toxicity and retrograde transmission. The resulting system requires four separate genes to be introduced into starter cells and the level of NS1 to be carefully controlled to suppress the undesirable properties of YFV. Also, it has not been demonstrated that YFV can trace circuits from Cre-positive neurons in a transgenic animal. Finally, the mechanism of transneuronal transfer of YFV is unknown, and it has not been shown to be strictly transsynaptic.

Herpes Simplex Virus (HSV) and Vesicular Stomatitis Virus (VSV) have also been used to trace neuronal circuits in the anterograde direction [3]. However, as with YFV, these viruses are toxic, travel retrogradely, and label second-order neurons that are not immediately downstream [7, 8]. Trans-Seq uses wheat germ agglutinin, a lectin, fused to mCherry (WGA-mCherry) to label cells transneuronally, allowing them to be sorted and subjected to single-cell transcriptome analysis [9]. This method does not allow for genetic access to postsynaptic cells and thus is not directly comparable to the techniques above or the one described in this paper. Furthermore, the mechanism by which transneuronal labeling occurs in all three of the above methods is unknown. Thus, there is no direct evidence that the three are strictly transsynaptic.

Here, we introduce ATLAS, a rationally designed protein capable of anterograde transsynaptic labeling, incorporating an antibody-like peptide, AMPA.FingR (AF), which binds to GluA1 but does not affect synaptic transmission. We provide evidence that ATLAS is released from presynaptic neurons, binds to AMPA receptors on the postsynaptic cell, and is taken up through endocytosis. We show in cultures that ATLAS can deliver Cre to postsynaptic neurons and mediate postsynaptic labeling through a floxed reporter both in culture and in vivo. ATLAS can be expressed explicitly in Cre-expressing neurons in transgenic animals and label neurons to which they project. In addition, it does not label neurons downstream of inhibitory neurons, providing evidence that labeling is exclusively transsynaptic. Finally, labeling is strictly anterograde, monosynaptic, and activity-dependent in vivo.

## RESULTS

### Generating AMPA.FingR, a binder to the N-terminus of GluA1

We used mRNA display, an in vitro selection procedure, to identify binders to a polypeptide encoding aa 1-394 of the extracellular N-terminus of Rat GluA1 produced in Sf9 cells (see methods). This bait was used to screen a library of mRNA-protein fusions encoding the 10FNIII Ig-like domain of human Fibronectin, with 17 random residues engineered into the BC and FG loops and a diversity of ∼10^13^ [10, 11]. After the 9th selection round, we cloned and sequenced 81 Fibronectin (Fn) clones, of which 24 were unique and full-length. We then put each Fn clone into an expression vector that placed a signal sequence from GluA1 at the N-terminus of the FN clone to target it to the secretory pathway and added a MYC tag for labeling. The resulting plasmids (ss-Fn-MYC) were co-transfected into COS7 cells with HA-GluA1 and Stargazin-V5. Surface labeling of MYC showed that four Fn clones labeled the surface of COS7 cells co-labeled with HA (Fig. S1). These four clones were expressed in dissociated cultures of cortical neurons. Of the four Fibronectins expressed in the cultures, one clone, Fn9.3, gave the most robust labeling (Fig. S1). We will call this clone AMPA.FingR (AF; FingR stands for **F**ibronectin **in**trabody **g**enerated with m**R**NA display).

We then generated AF-HA protein and found that it bound specifically to a band co-labeled with a GluA1 antibody on a western blot of neuronal lysate (Fig. 1a). To characterize AF further, we co-expressed it in dissociated cortical neurons with PSD-95.FingR-tagRFP, which binds to postsynaptic sites [10]. We found that AF labeling partially overlapped with PSD-95.FingR-tagRFP, consistent with a perisynaptic localization (Fig. 1b). AF also co-localized with endogenous Clathrin, consistent with localization to endocytic zones (Fig. 1c). Given that AF co-localizes with Clathrin and binds to GluA1, we tested whether AF-HA protein is endocytosed by adding AF-HA protein to the medium of cortical neurons in culture. We found that considerable AF-HA was detected in virtually all neurons labeled with MAP2 (37 of 38, n = 5 independent cultures) within 1 hour of addition (Fig. 1d). Furthermore, the addition of 10 mg/ml chlorpromazine, a clathrin inhibitor, significantly reduced the accumulation of AF-HA (4.2 ± 0.3 vs. 2.1 ± 0.5, p < 0.01, Mann-Whitney U test, Fig. 1e, f), confirming that internalization was mediated by endocytosis.

**Fig. 1.**
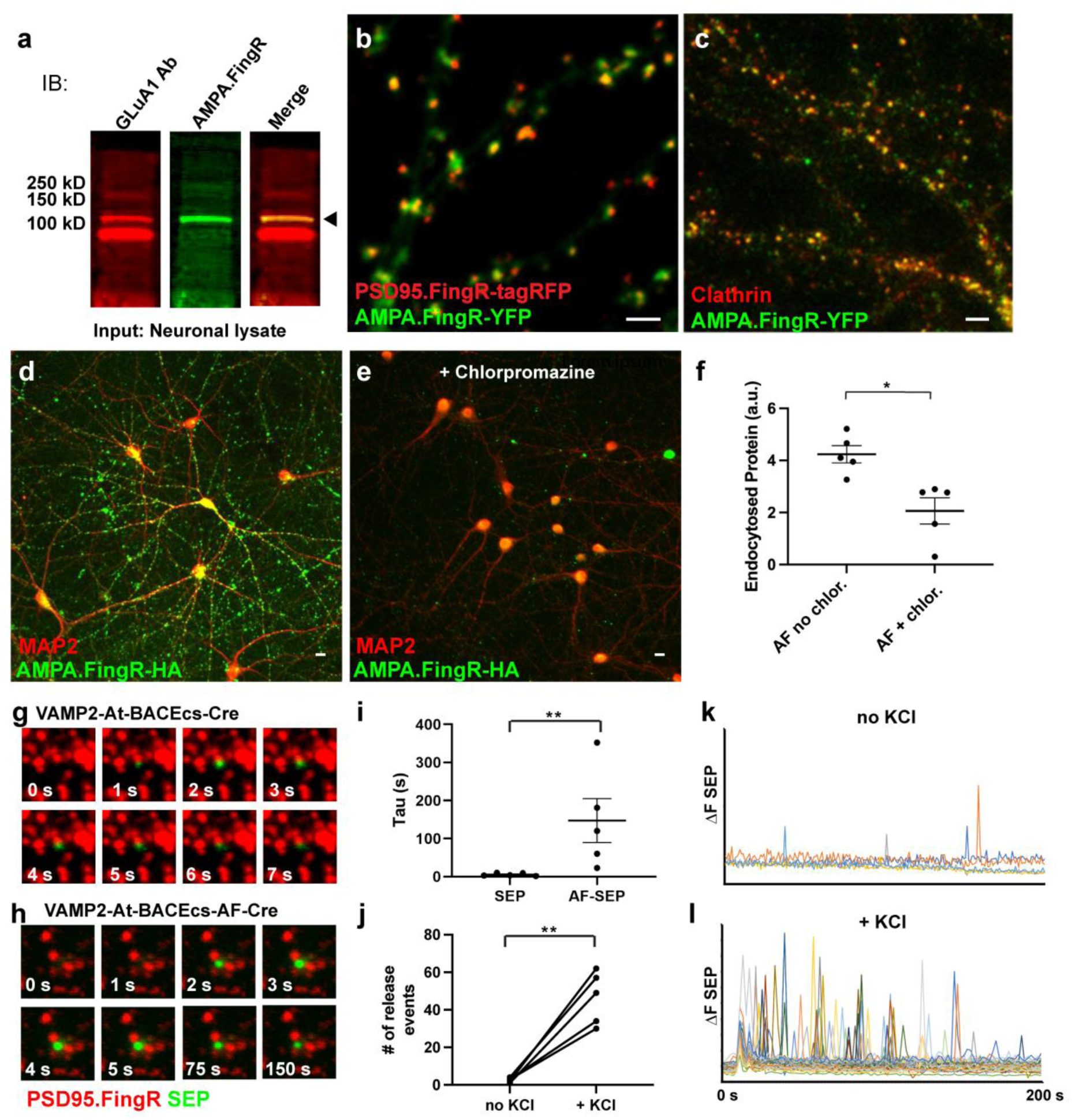
Cellular mechanisms of release and uptake by ATLAS. **a.** Western blot of lysate from dissociated neuronal cultures labeled with an anti-GluA1 Ab, AF-HA, and anti-GluA1 Ab + AF-HA (left to right). AF co-labels the higher MW band of two recognized by anti-GluA1 Ab. **b**. Live cortical neurons in culture expressing ss-AF-YFP (green) and PSD-95.FingR-tagRFP (red). AMPA.FingR appears to partially co-localize with PSD95.FingR, consistent with it localizing perisynaptically, adjacent to PSD-95. Scale bar 5 μm. **c**. Cortical neurons in culture expressing ss-AF-YFP fixed and co-labeled for endogenous Clathrin. Colocalization (yellow) is consistent with AF localization to endocytic zones. Scale bar 5 μm **d**. Cortical neurons in culture exposed to AF-HA (green) for 1 hour and co-labeled for endogenous MAP2 (red). Scale bar 25 μm. **e**. Same as d., but with the addition of 10 mg/ml chlorpromazine, a clathrin inhibitor, to the culture medium. Scale bar 25 μm. **f**. A significant reduction in endocytosed AF-HA protein with the addition of chlorpromazine. **g**. Release of SEP from presynaptic sites in cultured cortical neurons expressing VAMP2-At-BACEcs-SEP without additional KCl. PSD95.FingR-TagRFP (red), SEP (green). **h**. Same as g., but with VAMP2-At-BACEcs-AF-SEP. Note the increased brightness and longer duration of SEP fluorescence from VAMP2-At-BACEcs-AF-SEP vs. VAMP2-At-BACEcs-SEP, consistent with AF binding to the postsynaptic cell. **i**. τ of VAMP2-At-BACEcs-AF-SEP is significantly increased vs. VAMP2-At-BACEcs-SEP. **j**. Adding 100 mM KCl to the culture medium causes a dramatic increase in the frequency of release events of VAMP2-At-BACEcs-SEP. **k**., **l**. Graph of the differential of SEP fluorescence (ΔF) for VAMP2-At-BACEcs-AF-SEP with control medium vs. with medium containing 100 mM KCl. Each spike represents a single release event and does not reflect the duration of visible SEP fluorescence during each event (see Methods). * p < 0.05, ** p < 0.01. All tests Mann Whitney U test.

Given that AF likely binds to GluA1 receptors and is endocytosed postsynaptically, we reasoned that we could make a trans-synaptic tracer if we targeted AF to synaptic vesicles by fusing it to the luminal domain of the presynaptic protein, VAMP2 [12]. We inserted a β-Secretase cleavage site (BACEcs) [13] between AF and VAMP2 (to give VAMP2-BACEcs-AF), enabling endogenous BACE to cleave AF from VAMP2, leaving it free to exit the lumen of the synaptic vesicle upon exocytosis. In addition, an ALFA-tag (At) was added to label VAMP2 [14]. To test our strategy, we substituted superecliptic pHluorin (SEP) [12] for AF and co-expressed the resulting plasmid (VAMP2-At-BACEcs-SEP) with PSD-95.FingR-tagRFP in cortical neurons in culture. The release from acidic synaptic vesicles into the neutral pH of the synaptic cleft is expected to cause a sudden increase in the fluorescence of SEP, allowing it to be visualized [12]. Following overnight expression of VAMP2-At-BACEcs-SEP in neuronal cultures, we imaged SEP using fluorescence microscopy. We saw flashes of fluorescence lasting approximately 2-8 sec (images taken at 1 Hz) appearing at points opposite postsynaptic sites (Fig. 1g). This result is consistent with VAMP2 delivering SEP to synaptic vesicles, BACE cleaving off SEP, and SEP being released into the synaptic cleft. We also fused AF to SEP and expressed the resulting protein (VAMP2-At-BACEcs-AF-SEP) in culture and observed flashes that were much brighter and decayed with a much longer time constant (6 ± 2 sec vs. 147 ± 60 s, n = 5 independent cultures, Figs. 1h, i), a significant difference (P < 0.01, Mann Whitney U test). This result is consistent with AF binding to the postsynaptic membrane following release and remaining until it is endocytosed. To further test whether AF-SEP is released from synaptic vesicles, we examined whether depolarization increases the number of release events. We imaged the same cultures before and after adding 100 mM KCl and counted the number of bright flashes corresponding to release events. We found a ∼15-fold increase in release events with KCl (50 ± 6 vs. 3 ± 1 for control, P < 0.005, n = 5; Figs. 1j-l). Together, these results are consistent with AF-SEP being released from synaptic vesicles, moving across the synaptic cleft, binding to the adjacent postsynaptic neuron perisynaptically, and then being endocytosed.

Given that AF likely binds to the extracellular N-terminal domain of GluA1, we tested whether the expression of ss-AF-YFP affects synaptic transmission to hippocampal CA1 neurons. We found that in cells expressing ss-AF-YFP, the amplitudes of spontaneous EPSCs of AMPA and NMDA receptors and paired-pulse facilitation were not significantly different from control cells, consistent with the expression of AF not affecting synaptic transmission (Fig. s1).

### Transsynaptic tracing in vitro and in vivo

To generate a construct capable of mediating transsynaptic tracing, we replaced the SEP from the vector used in Fig. 1h with a recombinase, resulting in VAMP2-At-BACEcs-AF-Cre (ATLAS_Cre_). Given the above results, we expected that following expression of ATLAS_Cre_ in the presynaptic cell, VAMP2 would carry AF-Cre to synaptic vesicles, where it would be cleaved off and released into the synaptic cleft (Fig. 2a). AF-Cre should then bind to AMPA receptors at the postsynaptic membrane after which it should be endocytosed and carried to the nucleus. To facilitate this final step, we added the DNA binding domain from CCR5 [10, 15], which contains a robust nuclear localization signal, to the ATLAS protein. To test our strategy, we transfected ATLAS_Cre_ into dissociated cultures infected with an AAV carrying a floxed GFP (AAV8-DIO-GFP). We found cells with GFP and Cre that were not labeled with the starter cell marker, At, consistent with the transneuronal transport of Cre and recombination of the floxed allele, implying transport of Cre to the nucleus (Fig. 2b-e). Furthermore, when the presynaptic marker Synaptophysin-RFP was co-transfected with ATLAS_Cre_ in cultures infected AAV8-DIO-GFP, it co-localized with Cre in presynaptic puncta immediately adjacent to dendritic spines of GFP-expressing cells (Fig 2f). Thus, our results are consistent with ATLAS_Cre_ mediating transsynaptic tracing and providing genetic access to postsynaptic cells.

**Fig. 2.**
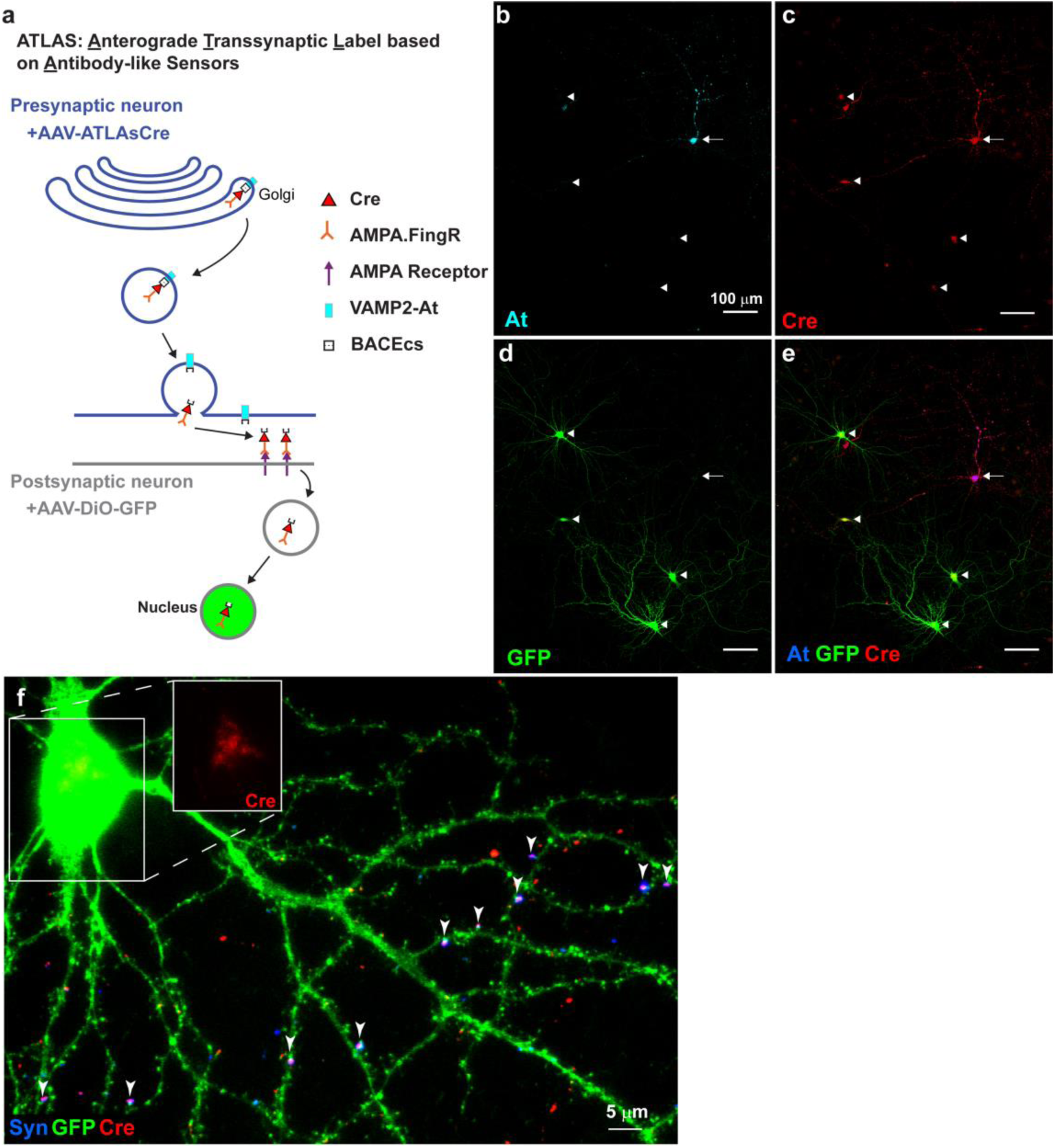
Design of ATLAS and testing in vitro. a. ATLAS_Cre_ protein (VAMP2-At-BACEcs-AF-Cre) is expressed in a presynaptic starter cell and targeted to synaptic vesicles by a VAMP2 domain. A BACE cleavage site within the ATLAS protein is cleaved before release. A peptide domain consisting of AMPA.FingR fused to Cre is released into the synaptic cleft and subsequently binds to the extracellular domain of postsynaptic GluA1. It is internalized through endocytosis and transported to the endosome. Finally, AMPA.FingR-Cre moves to the nucleus, where Cre can catalyze the expression of a floxed reporter gene. **b**. Cortical neuron in culture expressing ATLAS protein (arrow) following transfection, stained with anti-ALFA-tag nanobody (At). Note that four other neurons shown (arrowheads) do not express At. **c**. Cre staining of the same neurons as in b. including a neuron expressing ATLAS_Cre_ (arrow), and four not expressing ATLAS_Cre_ (arrowheads). Cre must have been transferred to the latter four neurons transneuronally. **d**. GFP labeling of postsynaptic cells (arrowheads, green) expressing Cre due to infection of AAV8-DIO-GFP. **e**. Combined labeling of At, GFP, and Cre showing presynaptic (arrow) and postsynaptic neurons (arrowheads). **f**. Cortical neuron from culture co-transfected with ATLAS_Cre_ and Synaptophysin-RFP and infected with AAV8-DIO-GFP. Co-localized Synaptophysin-RFP (blue) and Cre (red) labeling at presynaptic terminals are adjacent to dendritic spines of GFP-labeled postsynaptic neuron (green, arrowheads), consistent with an axon from a neuron expressing ATLAS_Cre_ synapsing onto the GFP-labeled postsynaptic cell. Diffuse Cre labeling is also present in the nucleus of the postsynaptic cell (inset), consistent with transsynaptic transport of Cre.

To test our strategy in vivo, we infected AAV8-ATLAS_cre_ into the medial prefrontal cortex (mPFC) and AAV8-DIO-mCherry into the striatum (Str) of an adult mouse (Fig. 3a). Following one week, we found At staining in mPFC, and abundant mCherry staining in the Str (Fig. 3b, c). In contrast, there was negligible mCherry staining in a control experiment without AAV8-ATLAS_cre_ (Fig. 3d). Because the projection from mPFC to Str is strictly anterograde [16], this result is consistent with AAV8-ATLAS_cre_ mediating the transsynaptic transport of Cre into the nucleus of the postsynaptic cell, causing mCherry labeling.

**Fig. 3.**
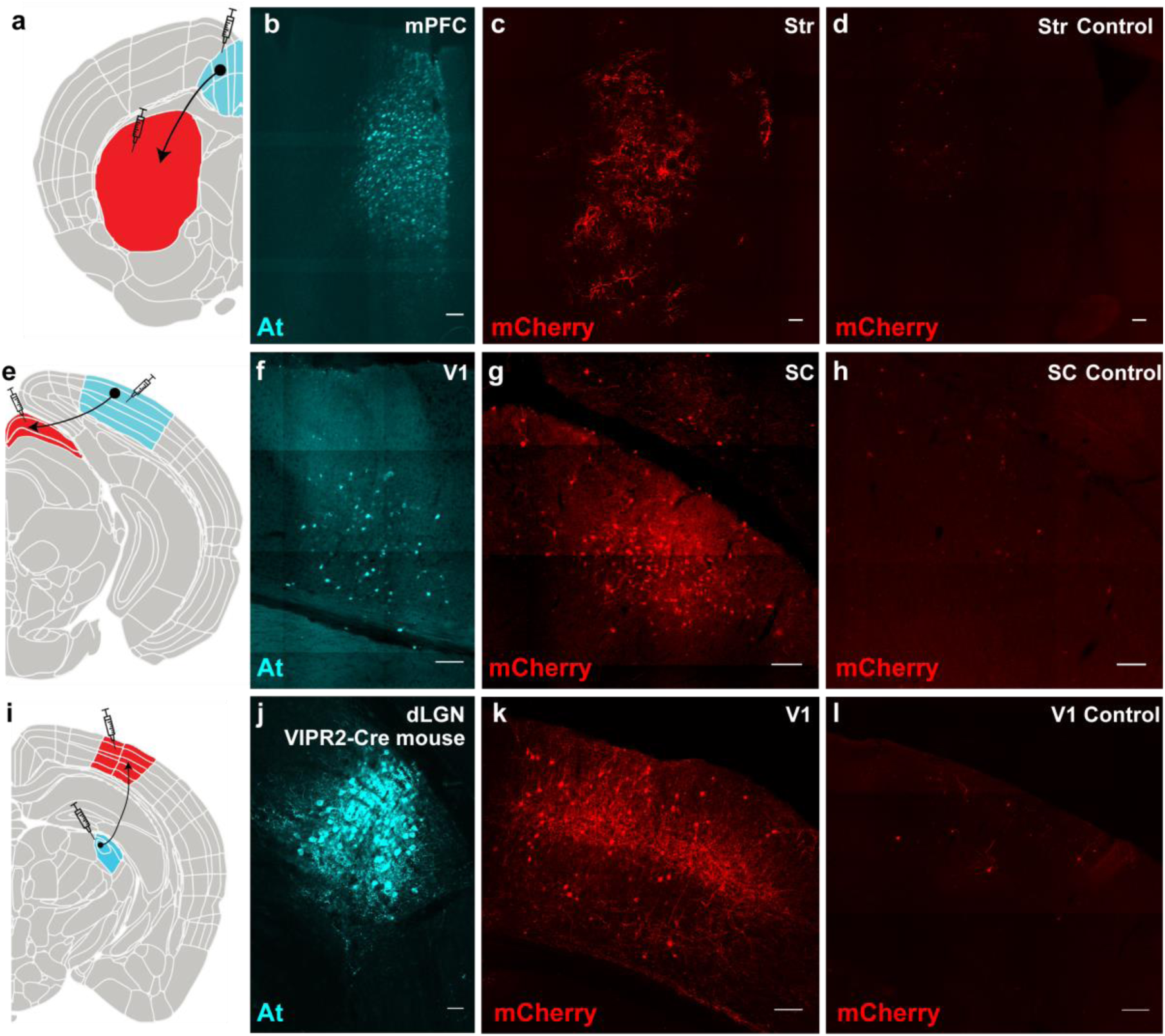
ATLAS mediates transsynaptic tracing in vivo. **a**. Schematic depicting injections of AAV8-ATLAS_Cre_ into the medial Prefrontal Cortex (mPFC, cyan) and AAV8-DIO-mCherry into the striatum (Str, red) to trace the unidirectional projection from the mPFC to the Str. **b**. ALFA-tag (At) staining of ATLAS_Cre_ in presynaptic cells in mPFC in a wild-type mouse. **c**. mCherry staining of postsynaptic cells in Str in the same brain as b. **d**. Control brain injected only with AAV8-DIO-mCherry in Str showing the extent of background labeling. **e**. Schematic depicting injections of DIO-AAV8-ATLAS_FLP_ and Lenti-Camkii-Cre into the primary visual cortex (V1, cyan) and AAV8-fDIO-mCherry into the Superior Colliculus (SC, red) to trace the unidirectional projection from V1 to SC. **f**. At staining of ATLAS_FLP_ in presynaptic cells in V1 in a wild-type mouse. **g**. mCherry staining of postsynaptic cells in SC in the same brain as f. **h**. Control brain injected only with AAV8-fDIO-mCherry in SC, showing the extent of background labeling. **i**. Schematic depicting injections of DIO-AAV8-ATLAS_FLP_ into the dorsal Lateral Geniculate Nucleus (dLGN, cyan) and AAV8-fDIO-mCherry into V1 (red) to trace the projection from dLGN to V1 in a VIPR2-Cre mouse. An advantage of this strategy is that ATLAS cannot be expressed in V1 even if the virus is transported retrogradely. **j**. At staining of ATLAS_FLP_ in presynaptic cells in dLGN. **k**. mCherry staining of postsynaptic cells in the V1 in the same brain as j. **l**. Control brain injected with only AAV8-fDIO-mCherry in V1 showing the extent of background labeling. Scale bars 100 μm.

### Tracing from genetically-determined cells

The ability to trace from genetically determined cell types is critical, given the heterogeneous distribution of different neuronal types in many brain regions. To mediate genetically determined tracing, we generated AAV8-DIO-ATLAS_FLP_, where expression of ATLAS is Cre-dependent and the payload is FLP (Flip recombinase). We tested this construct by tracing the projection from the primary visual cortex (V1) to the superior colliculus (SC), which is strictly anterograde [17]. We co-injected Lenti-Camkii-Cre (a lentivirus expressing Cre specifically in excitatory neurons) and AAV8-DIO-ATLAS_FLP_ into V1 and AAV8-fDIO-mCherry into SC. In addition to At staining in V1, we saw abundant staining for mCherry in SC compared to control (Fig. 3e-h). Note that an advantage of this strategy is that lentivirus does not have any intrinsic retrograde or anterograde transneuronal transport that could cloud the interpretation of the results, unlike virtually every pseudotype of AAV [4].

To test the ability of ATLAS to trace from labeled cells in transgenic Cre lines, we examined the projection from the dorsal Lateral Geniculate Nucleus (dLGN) to V1 in a VIPR2-Cre line expressing Cre in neurons in the dLGN [18]. We injected AAV8-DIO-ATLAS_FLP_ into dLGN and AAV8-fDIO-mCherry into V1. Staining of At was found specifically in the dLGN, and mCherry staining in V1 was dense in layer IV and less dense in layers II/III and V (Fig. 3i-l), consistent with known anatomy [19]. These results are consistent with ATLAS tracing anterogradely from Cre-expressing neurons in wild-type and transgenic mice.

### Labeling is transsynaptic

We showed that AF-Cre is secreted from presynaptic sites and endocytosed, likely through interaction with postsynaptic GluA1 (Fig. 1). Given the lack of other plausible mechanisms by which ATLAS can leave the presynaptic cell and enter the postsynaptic cell, it is likely that transneuronal labeling mediated by ATLAS occurs exclusively through synapses. To test this hypothesis in vivo, we asked whether transneuronal labeling could happen from inhibitory neurons. Given that uptake of ATLAS is mediated by GluA1 receptors, which are not found at inhibitory synapses, if we expressed ATLAS in inhibitory neurons, any transneuronal labeling would have to be nonsynaptic. Accordingly, we injected AAV8-DIO-ATLAS_FLP_ and fDIO-mCherry into the cortex of an SST-Cre mouse, where somatostatin-positive inhibitory interneurons express Cre [20]. When we examined the labeling in the cortex after two weeks, we saw no cells expressing mCherry that were not also labeled with At (Fig. 4a-c), indicating that no transneuronal labeling had occurred (Fig. 4a-c). Thus, the results of the above experiments, in combination with experiments in culture, lead us to conclude that ATLAS labels postsynaptic neurons exclusively through the transsynaptic transfer of proteins. Such an observation has not been made for any other transneuronal labeling method.

**Fig. 4.**
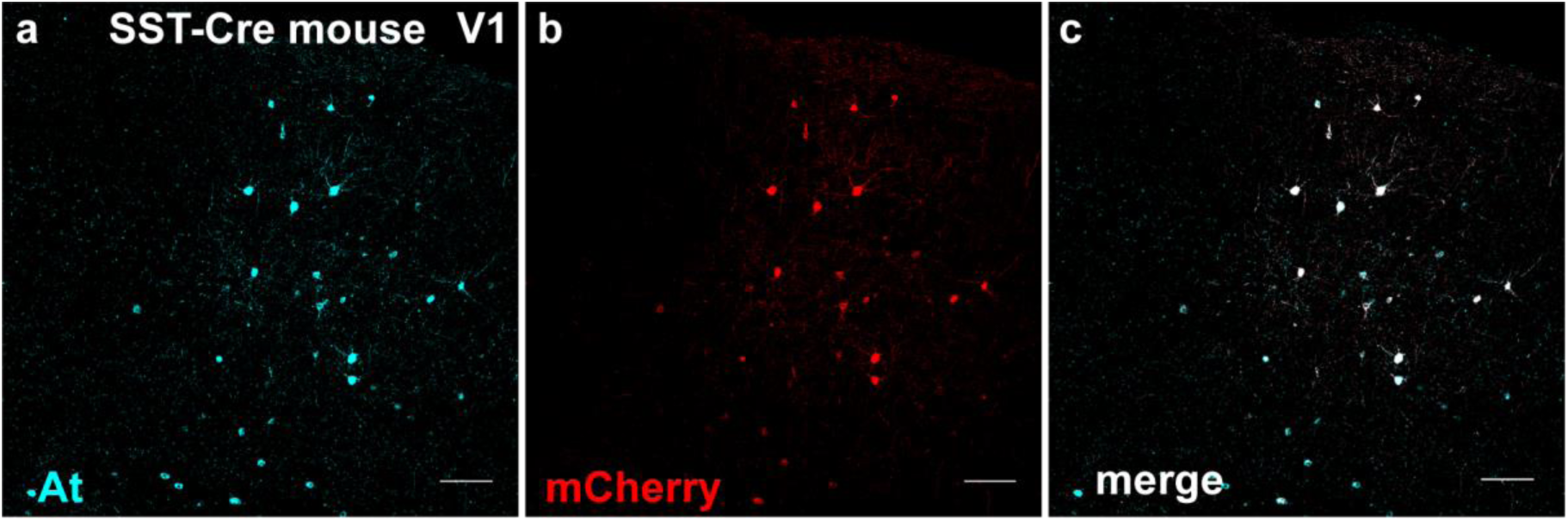
ATLAS does not mediate nonsynaptic transneuronal labeling. **a**. SST-Cre mouse with AAV8-DIO-ATLAS_FLP_ and AAV-fDIO-mCherry co-injected into V1. Labeling of inhibitory presynaptic neurons with At. **b**. mCherry-labeled neurons from the same brain as in a. **c**. Merged mCherry and At labeling shows that all neurons labeled with mCherry are also labeled with At, indicating that they are starter cells. Thus, there is no transneuronal labeling. Scale bar 100 μm.

### ATLAS is exclusively anterograde and monosynaptic

A significant limitation of many virally-based anterograde transneuronal labeling methods is that they also mediate labeling in the retrograde direction [4, 6]. Because AF binds specifically to the AMPA receptor and AF-Rec (AF-recombinase) is released from presynaptic terminals, it is unlikely to mediate retrograde transneuronal labeling (Fig. 1). To determine whether ATLAS mediates retrograde labeling in vivo, we injected AAV8-DIO-ATLAS_FLP_ and Lenti-hsyn-Cre into Str and AAV8-fDIO-mCherry into mPFC (Fig. 5a). Because the mPFC to Str projection is unidirectional and lentivirus does not spread retrogradely, any labeling in mPFC would be due to retrograde transneuronal labeling mediated by ATLAS. However, despite abundant At labeling in Str, indicating that ATLAS was expressed at a high level, we found no labeling above background in mPFC (Fig. 5b-d). These data are consistent with ATLAS not mediating retrograde transport.

**Fig. 5.**
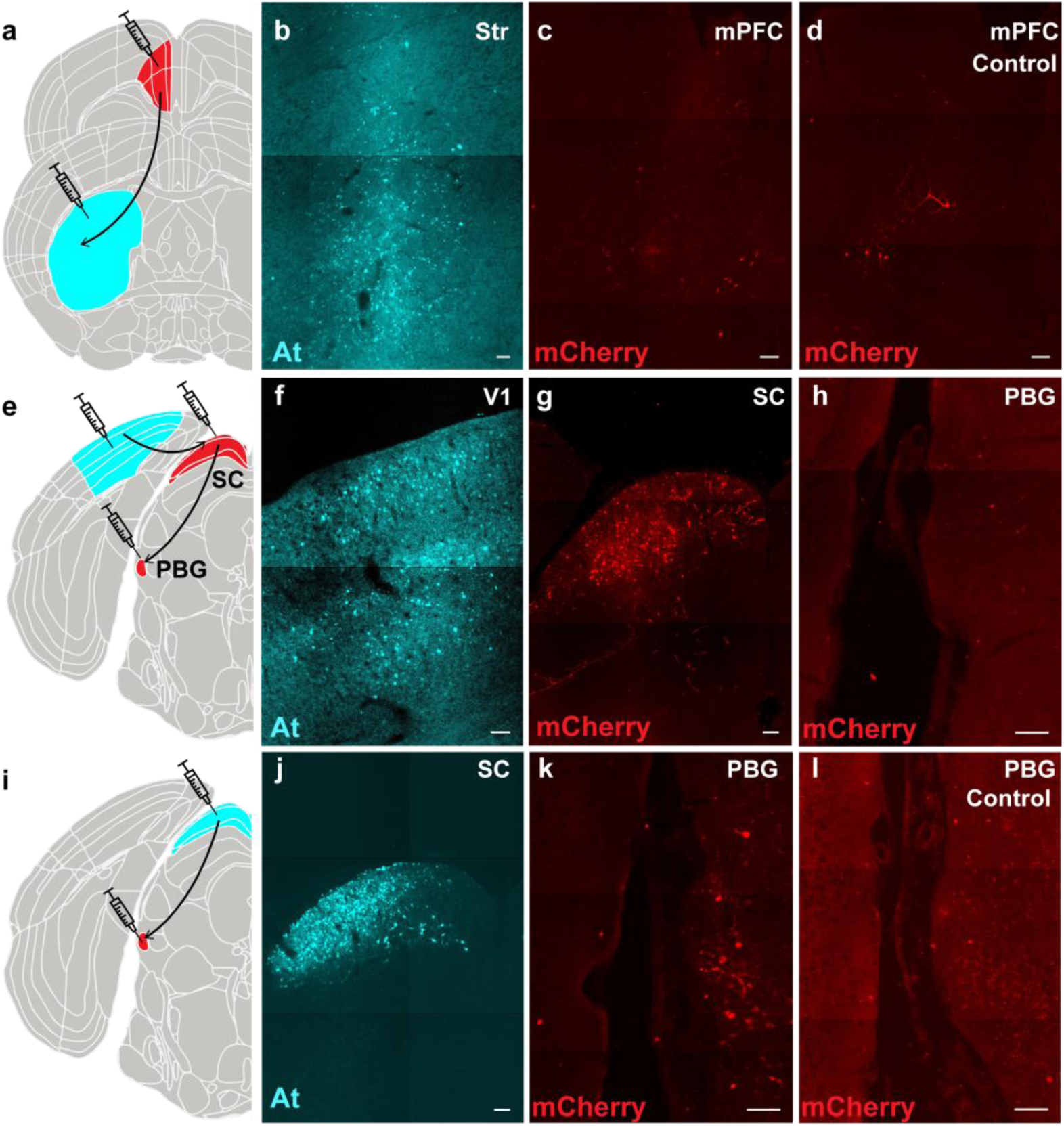
Transsynaptic labeling by ATLAS is exclusively anterograde and monosynaptic. **a**. Schematic of the projection from mPFC to Str showing injections of AAV8-DIO-ATLAS_FLP_ and Lenti-hsyn-Cre (Str) and AAV8-fDIO-mCherry (mPFC) for b, c. **b**. At labeling of cells in Str. **c**. mCherry labeling of cells in mPFC in the same brain as b. **d**. mCherry labeling of cells in mPFC in a control mouse brain with an injection of AAV8-fDIO-mCherry only into the mPFC. **e**. Schematic of the disynaptic pathway from V1 to SC to PBG showing injection of ATLAS_Cre_ (V1) and AAV8-DIO-mCherry (SC and PBG) for f-h. **f**. At labeling of presynaptic cells in V1. **g**. mCherry labeling of postsynaptic cells in SC from the same brain as f. **h**. mCherry labeling from the same brain as f. and g. shows shows no disynaptically labeled cells in PBG. **i**. Schematic of projection from SC to PBG showing injections of ATLAS_Cre_ (SC) and AAV8-DIO-mCherry (PBG) for j, k. **j**. At labeling of presynaptic cells in SC. **k**. mCherry labeling of postsynaptic cells in PBG from the same brain as j. **l**. mCherry labeling in PBG following injection of AAV8-DIO-mCherry only into PBG in a control mouse brain. Scale bar 100 μm.

To test whether ATLAS can mediate transsynaptic labeling that is monosynaptic, a critical property for transsynaptic tracing results to be interpretable, we tried tracing the polysynaptic pathway from V1 to SC to the Parabigeminal nucleus (PBG) [6]. We injected AAV8-ATLAS_Cre_ into V1 and AAV8-DIO-mCherry into SC and PBG (Fig. 5e). We found abundant transsynaptic mCherry labeling in SC but none in PBG (Fig. 5f-h). Furthermore, when we injected AAV8-ATLAS_Cre_ into Str, we saw abundant labeling in PBG, demonstrating that the Str to PBG pathway was intact (Fig. 5i-l). Thus, transsynaptic labeling mediated by ATLAS is exclusively anterograde and monosynaptic.

### Increasing efficiency of transsynaptic labeling through co-infection of BACE

To measure the efficiency of transsynaptic tracing by ATLAS, we injected AAV8-ATLAS_Cre_ in mPFC and AAV8-DIO-mCherry in Str, counted cells that we labeled after one week of incubation in four consecutive 25 μm thick sections, and calculated the density per mm^2^. Initially, we observed approximately 150 cells/mm^2^ labeled with mCherry in Str (150 ± 20 cells/mm^2^, n = 6, Fig. 6a, f). One possible inefficiency in our system is that there may be insufficient endogenous BACE in presynaptic cells to cleave AF-Cre from VAMP2. To test this hypothesis, we co-expressed exogenous BACE with ATLAS_Cre_ and measured the resulting efficiency. When we co-injected AAV8-BACE-HA and AAV8-ATLAS_Cre_ into mPFC in equal amounts, it did not affect the number of postsynaptic cells labeled with mCherry (Fig. 6b). However, when we injected AAV8-BACE-HA and AAV8-ATLAS_Cre_ in a 5:1 ratio, we saw a significant increase in the number of cells in Str labeled with mCherry (260 ± 10/mm^2^, n = 12; p < 0.001, Tukey’s multiple comparisons test, Fig. 6c-f). With this ratio, we saw approximately 10% of postsynaptic neurons labeled (Fig. 6d). Thus, when supplemented with exogenous BACE, ATLAS labels cells in the Str at a similar density to YFV in a similar experiment [6].

**Fig. 6.**
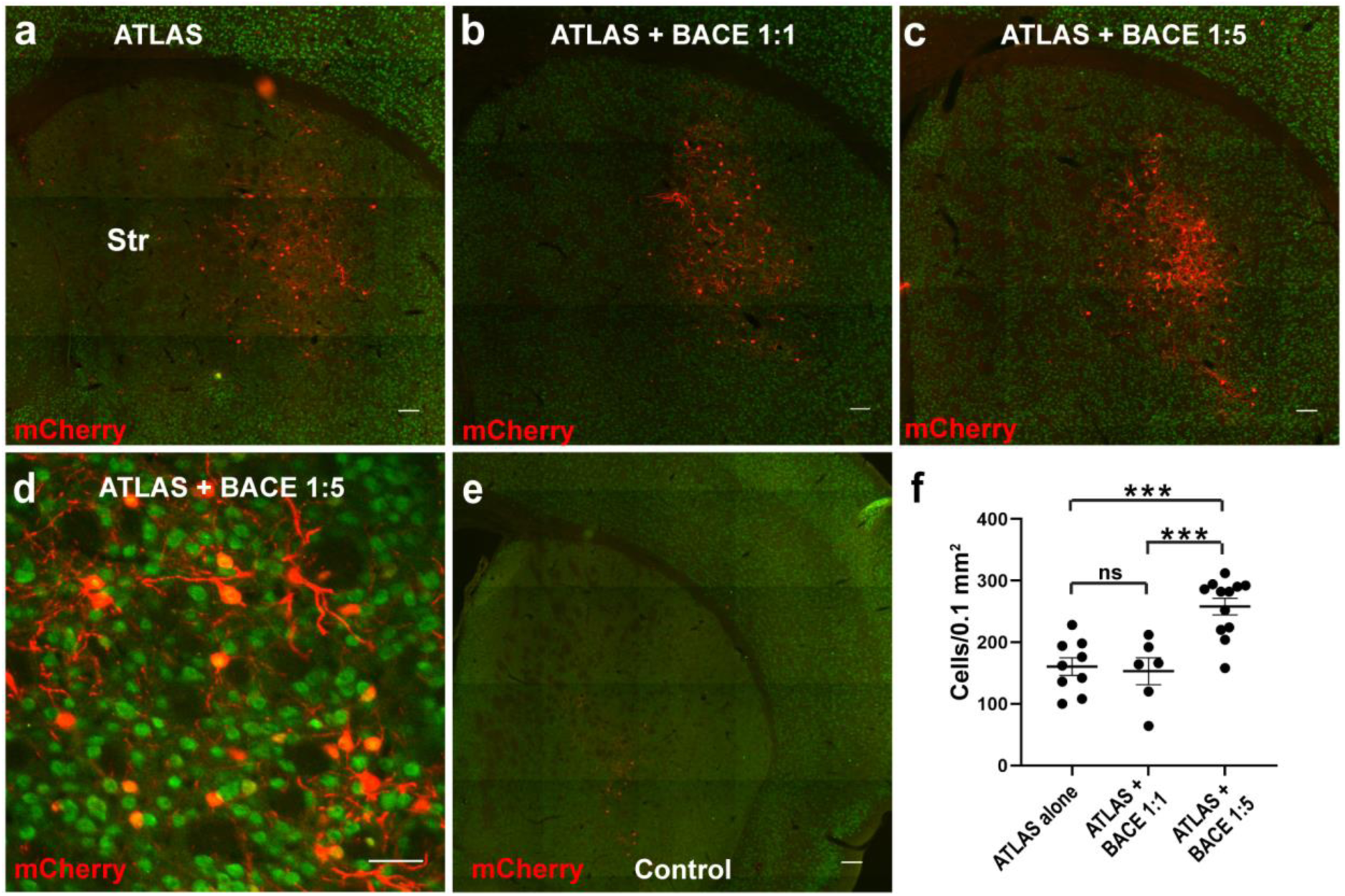
Improving the efficiency of ATLAS by co-expressing BACE. **a**. mCherry (red) and endogenous NeuN (green) labeling in Str in mouse brain infected with AAV8-ATLAS_Cre_ in mPFC and AAV8-DIO-mCherry in Str. **b**. Same as in a. with a 1:1 ratio of AAV8-ATLAS_Cre_:BACE-HA injected in mPFC and AAV8-DIO-mCherry in Str. **c**. Same as in a. with 1:5 ratio of AAV8-ATLAS_Cre_:BACE-HA injected in mPFC and AAV8-DIO-mCherry in the Str. **d**. Closeup of region in c. showing approximately 10% of all neurons labeled in Str. **e**. Control staining from an experiment with an injection of AAV8-DIO-mCherry only in Str. **f**. Quantitation shows that expressing ATLAS + BACE-HA 1:5 in mPFC leads to significantly higher labeling density in Str than ATLAS alone or ATLAS + BACE 1:1. *** p < 0.001 Tukey’s multiple comparisons test.

### Activity-dependence of ATLAS

Because ATLAS mediates transsynaptic labeling through the release of AF-Rec from synaptic vesicles and release is dramatically increased in high concentrations of KCl (Fig. 1j-l), transsynaptic labeling is likely activity-dependent. To test for activity dependence of transsynaptic labeling in vivo, we injected AAV8-ATLAS_Cre_ + AAV8- hHM3Dq-mCherry, an excitatory Dreadd [21], into mPFC and AAV8-DIO-GFP into Str (Fig 7a). We injected one group of mice with CNO intraperitoneally but not the other and compared GFP expression in Str (Fig. 7a-d). We found that GFP fluorescence intensity increased by 95% (37 ± 5 n = 4 vs. 19 ± 2 n = 4, Fig. 7g) in the mice injected with CNO vs. those without, a significant difference (p < 0.05, Mann Whitney U test, Fig. 7g), despite having roughly equal At expression in the mPFC (Fig. 7e, f). Thus, our experiments confirm that ATLAS mediates activity-dependent transsynaptic labeling.

**Fig. 7.**
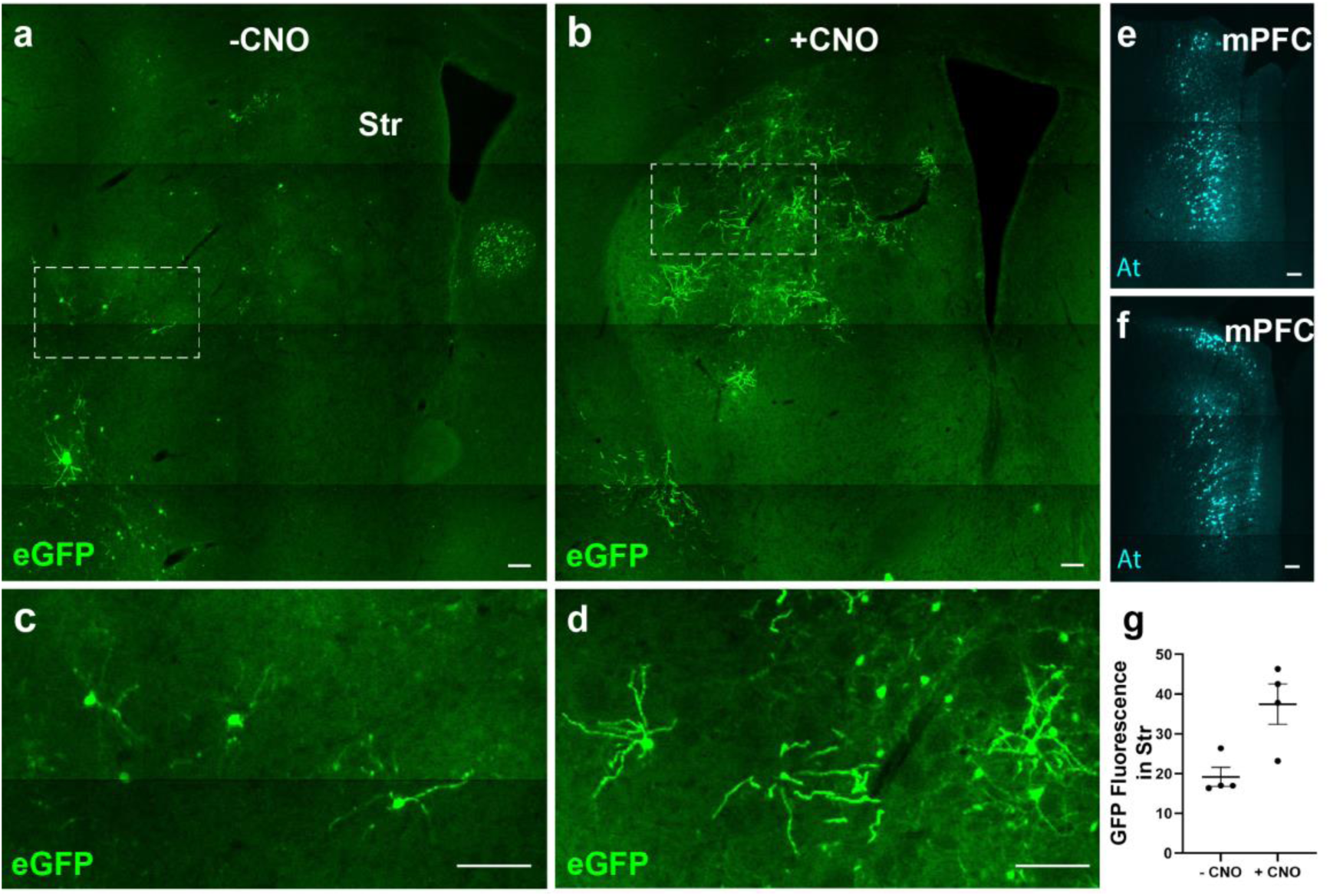
Transsynaptic labeling with ATLAS is activity-dependent. **a**. Str of mouse injected with AAV8-ATLAS_Cre_+ AAV8-hHM3Dq-mCherry in mPFC and AAV8-DIO-GFP in Str. **b**. Same as a, but with CNO injected intraperitoneally. **c**. Closeup of marked region in a. **d**. Closeup of marked region in b. **e**., **f**. Presynaptic staining of At in the mPFC showing similar expression of ATLAS_Cre_ for brains shown in a., b., respectively. **g**. Significantly higher GFP expression was measured in Str of mice injected with CNO than in control mice (* indicates p < 0.05, Mann Withney U test). All scale bars 100 μm.

## DISCUSSION

Here, we present ATLAS, a rationally designed protein for anterograde transsynaptic labeling based on a recombinant antibody-like protein, AF, which binds the N-terminal domain of GluA1. ATLAS differs from most other methods currently used for transneuronal labeling, which are based on naturally occurring viruses with gene deletions to eliminate undesirable qualities. ATLAS has advantages over modified viruses, including that its mechanism of action is precisely defined. Viruses such as rabies, yellow fever virus, AAVs, and the nonviral method Trans-Seq have modes of egress from the starter cells and entry into recipient cells that have not been precisely defined [5, 6, 9, 22]. In contrast, we have provided direct evidence that ATLAS releases a payload from synaptic vesicles at presynaptic sites and enters postsynaptic cells via binding to the extracellular domain of GluA1 followed by endocytosis of the receptor (Fig. 1). Another benefit of the nonviral nature of ATLAS is that, in principle, presynaptic cells do not have to be virally infected. Thus, it is possible to electroporate the ATLAS vector into presynaptic cells or an entire brain region in utero. In addition, it is possible to express ATLAS vectors presynaptically using non-toxic viruses such as Lentivirus or AAV, which enables ATLAS to mediate transsynaptic tracing without toxicity like that exhibited by rabies or yellow fever virus [6, 23].

Several qualities derive directly from the design of ATLAS, including that it is strictly anterograde. Its unidirectionality is due to the targeting of AF to synaptic vesicles, the specificity of AF binding, and the exclusively postsynaptic localization of its target, GluA1 (Fig. 1). We also designed ATLAS to mediate the release of AF-Cre from synaptic vesicles, which is activity-dependent. We further demonstrated that transsynaptic labeling with ATLAS is activity-dependent in vivo (Fig. 7). In addition, because ATLAS works via a protein expressed in presynaptic cells, it was easy to make tracing from presynaptic cells Cre-dependent by making expression of ATLAS_FLP_ Cre- dependent (Fig. 3). Thus, ATLAS can be used to trace circuits from genetically specified cells, such as Cre-expressing cells in transgenic mice, which has not been demonstrated using other anterograde tracing methods. Another way to express ATLAS in specific cell types is through Cre expression driven by a cell-type specific promoter or of cell-type specific expression of ATLAS itself. Also, there is no mechanism by which ATLAS can mediate retrograde or polysynaptic labeling, resulting in strictly monosynaptic anterograde labeling, a property of ATLAS that we demonstrated in vivo (Fig. 5). Finally, we used the fact that ATLAS works only at excitatory synapses to show it is incapable of nonsynaptic transfer by observing that it did not undergo transneuronal labeling when expressed in inhibitory neurons in vivo (Fig. 4).

Because ATLAS does not have an intrinsic amplification mechanism like rabies and similar viruses [24], it requires the expression of a recombinase-dependent reporter in the postsynaptic cell. While this could be interpreted as a drawback, it is also a benefit because amplification, such as in rabies or VSV infection, is associated with toxicity [25, 26]. In the case of AAV1, which also requires amplification, a transgenic mouse with a floxed reporter transgene can be used instead of postsynaptic injections [4, 5]. Unfortunately, efficient and consistent transsynaptic labeling did not occur when we expressed ATLAS_Cre_ in a floxed reporter mouse (data not shown). Although the reason for this is unknown, it is possible that the presence of the virus somehow facilitates the release of the AF-Rec from the endosome and/or transport to the cytoplasm and nucleus in the postsynaptic cell. Previously, these processes were found to be limiting for protein transduction [27, 28]. The requirement for a postsynaptic virus means that postsynaptic target areas must be identified a priori, which is not required for most virus-based transneuronal labeling techniques. However, this should not be a difficult problem to surmount as regions postsynaptic to specific cell types are easily identified by exogenously expressing a presynaptic marker such as VAMP2 [29].

Nonetheless, ATLAS would be improved if postsynaptic targets could be labeled globally without injecting a second virus. One possible partial solution to this problem would be to use a systemic injection of a virus such as PHP.eb [30, 31] in combination with an injection of the ATLAS presynaptic vector. Although such a method would still require administering a second virus, it would not require knowledge of the postsynaptic region or a second stereotaxic injection. A single injection could also provide broad coverage of many different brain regions.

Perhaps the most significant advantage of a rationally designed tracer such as ATLAS over traditional viral tracers is that its modular components can be modified and optimized. In this way, it will be possible to evolve ATLAS to change its properties and improve performance in a manner similar to sensors such as GCaMP [32] or GRAB sensors [33]. For instance, the form of ATLAS described in this paper works only for anterograde tracing from excitatory neurons. However, if the AF were replaced with a recombinant binder to an inhibitory postsynaptic receptor, such as the GABA receptor, it could, in principle, be used for anterograde tracing from inhibitory neurons. Similarly, the substitution of AF for binders of other neurotransmitter receptors such as acetylcholine, or even neuromodulators such as dopamine or serotonin could be the basis of other transneuronal tracing systems, which is an exciting prospect given the proliferation of recombinant postsynaptic receptor binders [34]. Another possibility is to use activity dependence of ATLAS labeling to mark circuits that respond to specific stimuli. This technique would be especially useful if ATLAS tracing could be gated to occur during a particular time interval. Finally, it may be possible to modify ATLAS to label circuit changes during events such as memory formation [35] or specific developmental processes such as synaptic pruning [36].

## ACKNOWLEDGEMENTS

We thank all Arnold lab members for helpful discussions. We also thank Alexandra Delgadillo, Melissa Qin, Heesung Sohn, Aida Bareghamyan, and James Qi for technical assistance and Fan Wang for crucial input at the beginning stages of this project. We thank Emily Liman for their insightful comments on the manuscript. This work was supported by a grant (MH116989) to D.A. from NIMH and the Brain Initiative.

## AUTHOR CONTRIBUTIONS

D.A. conceived and designed the study, assisted with data analysis, and wrote the paper with input from all authors. J.R. generated the AMPA.FingR, performed the biochemistry and neuronal culture-related experiments, and designed and generated the plasmid constructs. W.W. and H.H. performed and analyzed all in vivo tracing experiments. J.R., W.W., and H.H. generated the AAVs and lentiviruses. B.H. and S.R. designed and S.R. performed the electrophysiology experiments.

## Methods

### Animal preparation and stereotaxic surgery

All experiments were performed in accordance with NIH Guidelines for the Care and Use of Laboratory Animals. All procedures used in this study were approved by the Animal Care and Use Committee at the University of Southern California. Male and female Wild-type (C57BL/6J, Jackson Laboratories, stain #000664), Vipr2-Cre (Cre expression in dLGN, Jackson Laboratories, strain #031332), and Sst-IRES-Cre (Cre expression in somatostatin interneurons, Jackson Laboratories strain #013044) mice aged 2–6 months were used in this study. Mice were housed in a light-controlled (12 h light/dark cycle) environment with ad libitum access to food and water. For stereotaxic viral injection, mice were anesthetized initially in an induction chamber containing 4% isoflurane mixed with oxygen and then transferred to a stereotaxic frame equipped with a heating pad. Anesthesia was maintained throughout the procedure using continuous delivery of 1.5% isoflurane through a nose cone at a rate of 0.5 L/min. The scalp was shaved, and a small incision was made along the midline to expose the skull. After leveling the head relative to the stereotaxic frame, injection coordinates based on Paxinos and Franklin’s 4th Edition [37] were used to mark the location on the skull directly above the target area, and a small hole (0.5 mm diameter) was drilled. Viruses were delivered through pulled glass micropipettes using pressure injection via a micropump. Total injection volumes ranged from 50 to 600 nl, at 20 to 60 nl/min. Following injection, the micropipette was left in place for 15 min to minimize virus diffusion into the pipette track. After withdrawing the micropipette, the scalp was sutured closed. Animals were recovered from anesthesia on a heating pad to minimize inflammation and discomfort and returned to their home cage. Animals were administered Ketofen (5mg/kg) at the beginning of the surgical procedure and again every 24 h for 2d following surgery. Bregma and Lambda coordinates are listed in millimeters as follows: AP is the distance from Lambda, ML is the distance from midline, and DV is the distance from Lambda except for V1 and SC, which use the distance from the surface of the cortex. Medial Prefrontal Cortex (mPFC) (6.3, 0.3, 1.7); Str (5.1, 1.8, 3.4); dorsal lateral geniculate nucleus (dLGN) (2.2, 2.2, 2.9); Primary Visual cortex (V1) (0.8, 2.4, 0.6); Superior colliculus (SC) (0.6, 0.8, 1.2); Parabigeminal nucleus (PBG) (0.5, 1.8, 3.3);

### Histology

Mouse brains were perfused with 40mL of phosphate-buffered saline (PBS) followed by 40mL of 4% paraformaldehyde in PBS. The brains were then fixed overnight in 4% paraformaldehyde at 4°C and dehydrated in 30% sucrose solution for at least 48 hours. The brains were embedded in tissue freezing medium at -45 °C and then sectioned to a thickness of 25um-35um using a cryostat. For immunohistochemistry, the brain sections were washed in PBS and incubated overnight in primary antibodies diluted in PBS containing 4% goat normal serum and 0.5% triton X-100. The primary antibodies used are as the following: (animal) anti-RFP, Rockland (600-401-379), 1:1000; (animal) Anti-GFP antibody, AVES (gfp-1020), 1:1000; 1:1000; Fluotag-X2 anti-ALFA, NanoTag (N1502-ALFA647-L), 1:1000 The sections were then washed with PBS and incubated in corresponding secondary antibodies for 2 hours. After incubation, the sections were washed with PBS and mounted onto glass slides with DAPI Flouromount-G, EMS. The secondary antibodies used in this study and their dilution are as follows: Alexa Flour 488-conjugated goat anti-chicken or anti-mouse IgG, Thermo Fisher, 1:1000; Alexa Flour 594-conjugated anti-rabbit IgG, Thermo Fisher, 1:1000; Alexa Flour 647-conjugated goat anti-mouse IgG, Thermo Fisher, 1:1000

### Electrophysiology

All experiments were performed in accordance with NIH Guidelines for the Care and Use of Laboratory Animals, and all procedures were approved by the Institutional Animal Care and Use Committee of the University of Southern California. As previously described, 400 μm rat organotypic hippocampal slice cultures were prepared from male and female P6 to P8 Sprague Dawley rats [38]. Tissue was isolated, and an MX-TS tissue slicer (Siskiyou) was used to make 400 μm transverse sections. Tissue slices were placed on squares of Biopore Membrane Filter Roll (Millipore) and placed on Millicell Cell Culture inserts (Millipore) in 35 mm dishes. Slices were fed 1mL of culture media containing MEM + HEPES (Gibco Cat#12360-038), horse serum (25%), HBSS (25%) and L-glutamine (1 mM). Media was exchanged every other day. Whole-cell recordings were performed on DIV7-8. During recordings, slices were maintained in room-temperature artificial cerebrospinal fluid (aCSF) containing 119 mM NaCl, 2.5 mM KCl, 1 mM NaH_2_PO_4_, 26.2 mM NaHCO_3_ 11 mM glucose, 4 mM CaCl2, and 4 mM MgSO4. 5 μM 2-chloroadenosine and 0.1 mM picrotoxin were also added to the aCSF to dampen epileptiform activity and block GABA_A_ receptor activity, respectively. Osmolarity was adjusted to 310-315 mOsm. aCSF was saturated with 95% O2/5% CO2 throughout the recording. Borosilicate recording electrodes were filled with an internal whole-cell recording solution containing 135 mM CsMeSO_4_, 8 mM NaCl, 10 mM Hepes, 0.3 mM EGTA, 5 mM QX-314, 4 mM Mg-ATP, and 0.3 mM Na-GTP. Osmolarity was adjusted to 290–298 mOsm and pH-buffered at 7.3–7.4.

CA1 pyramidal neurons were identified using differential interference phase contrast microscopy while GFP-expressing transfected neurons were identified using epifluorescence microscopy. Dual whole-cell recordings of CA1 pyramidal neurons were made through simultaneous recordings from a transfected neuron and a neighboring, non-transfected control neuron. Synaptic responses were evoked by stimulating with a monopolar glass electrode filled with aCSF in the stratum radiatum. Membrane holding current, pipette series resistance and input resistance were monitored throughout recording sessions. Data were acquired using a Multiclamp 700B amplifier (Molecular Devices), filtered at 2 kHz, and digitized at 10 kHz. AMPAR-evoked EPSCS (-eEPSCs) were measured at -70mV. NMDAR-eEPSCs were measured at +40 mV and were temporally isolated by measuring amplitudes 150 ms following the stimulus, at which point the AMPAR-eEPSC had completely decayed. For paired-pulse facilitation, paired pulses at 40ms inter-pulse interval were delivered, and the ratio of peak amplitudes was analyzed to generate barplots depicting mean ± SEM. Data analysis was performed using Igor Pro (Wavemetrics). In the scatter plots for simultaneous dual whole-cell recordings, each open circle represents one paired recording, and the closed circle represents the average of all paired recordings. No more than one paired recording was performed on any given hippocampal slice.

#### Biolistic Transfection

Sparse biolistic transfections were performed on day in vitro 1 (DIV1) as previously described [39, 40]. 50 μg of mixed plasmid DNA encoding ss-AF-YFP was coated on 1μm-diameter gold particles in 0.5 mM spermidine, precipitated with 0.1 mM CaCl_2_, and washed four times in pure ethanol. These DNA-coated gold particles were then coated onto PVC tubing, dried briefly using ultra-pure N_2_ gas, and stored at 4 °C in desiccant. Before use, the gold particles were brought to room temperature and delivered to slice cultures via a Helios Gene Gun (BioRad). Construct expression was confirmed by YFP epifluorescence.

#### Target preparation, mRNA display, and screening

The amino terminal domain of GluA1 (1-394 a.a.) fused to a biotin acceptor tag (AviTag, Avidity) grown in suspension cultures of Sf9 cells was used as a target for the mRNA display selection. mRNA display procedure was carried out as described in [41]. Screening for binding FingRs was performed in Cos7 cells expressing ss-GluA1-HA (where ss is the signal sequence of GluA1) and Stargazin-V5 with the candidate FingRs fused to a Myc tag. Cos7 cells were initially stained live with anti-MYC antibody and then fixed, permeabilized, and stained for HA and V5. Fns labeling the surface of cells expressing GluA1 and Stargazin were then expressed in dissociated cultures and tested for the ability to label endogenous GluA1.

#### Dissociated cultures

Pups from timed pregnant Sprague Dawley rats (E19) (Charles River) were used for dissociated cultured cortical neurons as described in [42]. Briefly, cortices from E19 pups were trypsinized in Neuronal Isolation Enzyme with papain (ThermoScientific 88285) and triturated into a single cell suspension. Cells were plated at a density of 1.5 × 10^5 per well on 6-well plates in Neurobasal media (Gibco 21103-049) containing B27 supplement (Gibco 17504-044), Fetal Bovine Serum Characterized (Cytiva SH30071.03), glutaMAX-I (Gibco 2523105) and gentamicin (Sigma G1397). Media was changed after 4 hours, and after 7 days, it was replaced with neurobasal media without Fetal Bovine Serum.

#### Transfection

Cultured neurons were transfected using the CalPhos Mammalian Transfection Kit (Takara Cat# 631312). Briefly, 8 ug of plasmid DNA was incubated with sterile H2O and 2M CaCl_2_ to 100 ml; this mixture was added to 100 ml of 2X HEPES-buffered saline and incubated at room temperature for 20 minutes. The mixture was then added to the cultured neurons dropwise and incubated for 1-3 hours for crystal formation. After incubation, the media was removed, and neurons were washed with Hepes-NaCl buffer (140mM NaCl, 5mM KCl, 24mM D-glucose, and 10 mM HEPES, pH 7.2). After washing, transfected neurons were maintained in conditioned neurobasal media.

#### Immunocytochemistry

Dissociated cultured neurons were fixed in 4% Paraformaldehyde (EMS 15714) for five minutes, followed by three, 5 minute washes in 1x phosphate buffered saline (PBS). After washes, neurons were blocked for 30 minutes in blocking buffer (5% Normal goat serum, 1% Bovine serum albumin, 0.1% Triton-100 in PBS). Following the blocking step, neurons were incubated with primary antibodies for 1 hour at room temperature (anti-GFP, Aves GFP-1020; anti-Map2, Abcam ab5392; anti-clathrin, GeneTex GTX80218; anti-mCherry, Rockland M11217; anti-RFP, Invitrogen; anti-GluA1, EMD Millipore ABN241; anti-Psd95, NeuroMab 75-348; anti-HA, cell signaling 3724S; anti-Cre, EMD Millipore MAB3120), followed by three, 5 minute washes in PBS. Neurons were incubated in secondary antibodies in blocking buffer for 1 hour at room temperature in the dark (Alexa Fluor 488 goat anti-chicken, Invitrogen A-11006; Alexa Fluor 594 goat anti-rabbit; Invitrogen A-11012; Alexa Fluor 594 goat anti-mouse Invitrogen A-11005, Alexa Fluor 647 goat anti-mouse Invitrogen A-21235; Alexa Fluor 647 goat anti-rabbit, Invitrogen A-21244; JF549 halotag ligand, Janelia; JF647 halotag ligand, Janelia; FluoTag-X2 anti-alfa 647, NanoTag biotechnologies N1502 AF647, FluoTag-X2 anti-alfa 568, NanoTag biotechnologies, N1502 AF568). Neurons were mounted using Dapi-fluoromount-G (Electron microscopy services 17984-24).

#### Microscopy

Fixed dissociated neurons were imaged using Olympus IX-71 and IX-81 microscopes. Confocal images were acquired using a Zeiss 880 microscope.

#### Western blot

Lysates from cultured neurons were collected in NP40 lysis buffer (150 mM NaCl, 100 mM Tris-HCl, 1% NP40 in PBS) containing 1X protease inhibitors (cOmplete ULTRA tablets, Roche, cat# 05892970001). Samples were run on mini-protean TGX gels (Biorad, cat# 4569034) and transferred to LF PVDF membrane (Biorad, cat# 1620264). The membrane was blocked in 5% dry milk in PBS. The membrane was probed with anti-GluR1, EMD Millipore ABN241; anti-HA, cell signaling 3724S; anti-tubulin, Sigma T6168; and purified AF-HA. Membranes were imaged with the LI-COR Odyssey IR imaging system.

#### Protein purification

AF-HA protein was purified from the media of HEK293T cells. Briefly, HEK293T cells were transfected with the plasmid pCAG-ss-AF-3XHA-TEVcs-HALOTag using Lipofectamine 2000 (Invitrogen, Cat# 11668-019) following the manufacturer’s protocol. The medium was collected from transfected HEK293T cells containing the secreted protein AF-HA-TEVcs-HALOTag. The medium was incubated with HaloLink resin (Promega, Cat# G1914) overnight at 4 °C. The resin was washed 3 times with wash buffer (100 mM Tris, pH7.6, 150 mM NaCl, 0.5% Triton-X), and the resin was then incubated with TEV protease (New England Biolabs, Cat# P8112S) overnight at 4° C. Next, the supernatant was collected and incubated with NEBExpress Ni Resin (New England Biolabs, Cat# S1428S) overnight at 4° C to remove the TEV protease from the purified protein. Purified protein was then dialyzed into 20mM HEPES, 250mM NaCl pH 7.4, and concentrated using the Amicon Ultra centrifugal filters (cat# UFC501096).

#### Endocytosis experiments

Neuronal cultures were incubated with 1 μM purified AF-HA protein for 1hr at 37 °C. Next, cultures were washed 3 times with 1X Hanks Balanced Salt Solution containing 10 mM HEPES, fixed with 4% paraformaldehyde, and stained for anti-HA (Cell signaling 3724S) and anti-Map2 (Abcam ab5392).

#### Presynaptic release experiments

Plasmids encoding VAMP2-BACEcs-SEP (1ug) or VAMP2-BACEcs-AF-SEP (1 μg) with HA-BACE (2 μg) were transfected into 15-18 DIV cultured neurons using the calcium phosphate method (see below). Two days before transfection, the cultures were infected with AAV8-Cre and AAV8-PSD95.FingR-TdTomato to label postsynaptic sites. Neurons were imaged 3 to 5 days later with or without 100 mM KCl. Time-lapse imaging was performed for 200 frames at an interval of 1000 ms and an exposure of 500 ms. Time constants were measured using curve fitting functions in GraphPad. Videos of ΔF of SEP during release events were generated using ImageJ by taking the difference of adjacent image frames to eliminate background and SEP fluorescence beyond the initial frame after release.

#### Transsynaptic tracing in culture

Neuronal cultures (14 DIV) were transfected with plasmids for ATLAS (1 μg) and HA-BACE (2 μg) using the calcium phosphate method. Transfection was followed by infection with AAV8-DIO-GFP (with human synapsin, hSyn, promoter) and incubated for 7 days. Neurons were fixed and probed as described.

#### Synaptophysin-RFP

Plasmids for ss-AF-YFP (200 ng) and Synaptophysin-RFP (200 ng) were transfected into 13 DIV cultured cortical neurons and imaged live the next day

#### AAV virus production and purification

Briefly, one day before transfection, Hek293T cells were plated at 1.5 × 10^7^ cells per 15 cm dish. Plasmids for pHelper, AAV2/8 (Gifts from Dr. Fan Wang), and gene of interest were transfected using PEI (1mg/ml). The cells were cultured for 72 hrs. Cells were scraped, and the pellet from 6 dishes was resuspended in 4 ml of cell lysis buffer (150 mM NaCl, 50 mM Tris-HCl, pH 8.5). The pellet was then frozen in a dry ice/ethanol bath and thawed in a 37° C water bath for three cycles. The cell lysate was thawed in a 37 °C water bath, and benzonase was added at a final concentration of 50 U/ml for 30 min. This was followed by centrifugation at 4,500 rpm for 30 min at 4 °C in a tabletop centrifuge. Subsequently, vector-containing supernatant was collected and loaded over an iodixanol density gradient, centrifuged at 50,000 rpm for 2 h at 18 °C, and concentrated using VIVASPIN 20 10K MWCO filter tubes.

#### Lentivirus production and purification

Briefly, one day before transfection, Hek293T cells were plated at 1.5 × 10^7^ cells per 15 cm dish. Plasmids for pMD2.G (Addgene #12259), psPAX2 (Addgene #12260), and gene of interest were transfected using PEI (1mg/ml) and cultured for 72 hrs. Cell supernatant was collected and centrifuged at 1,500g for 15 min. The virus-containing supernatant was filtered through a 0.45 μm filter flask, loaded into sterilized 50 ml centrifuge tubes with 7.5 ml of 10% sucrose solution, and centrifuged for 4 hours at 10,000g and 4 °C. After centrifugation, the supernatant was 50 μl of ice-cold PBS added (without Ca^2+^ and Mg^2+^) to cover the virus pellet and let sit at 4 °C for 2 hours. The pellet was resuspended gently and spun at 7,000 rpm for 5 min. Finally, the supernatant was concentrated using VIVASPIN 20 30K MWCO (SARTORIUS) filter units with PBS.

#### Plasmid Constructs

The signal sequence from GluA1 (AA 1-18) was inserted upstream of the AMPA.FingR sequence and YFP was inserted downstream of the AF sequence to generate **ss-AF-YFP**. For all constructs containing VAMP2, the complete sequence of VAMP2 was inserted upstream on an ALFA Tag (SRLEEELRRRLTE) followed by a BACE (β-site Amyloid Precursor Protein-cleaving enzyme 1) cut site (EISEVNLDAEFR). In the case of **VAMP2-BACEcs-SEP**, the BACE cleavage site (BACEcs) was followed by super ecliptic pHlurion (SEP). For **VAMP2-BACEcs-AF-SEP**, the BACEcs was followed by the AF sequence and SEP downstream of AF. For **DIO-ATLAS_FLP_**, Cre was removed from the ATLAS construct and replaced with a Myc Tag (EQKLISEEDL) followed by the recombinase Flp. Nested Lox sites were then added on either side of the coding sequence. GFP-HA-BACE1 plasmid was purchased from addgene (#165032), and the GFP was removed to make **HA-BACE**. AAV constructs were generated by ligating the above-mentioned sequences into a pAAV plasmid containing the human synapsin (hsyn) promoter and a woodchuck hepatitis virus post-transcriptional regulatory element (WPRE). The plasmid Lenti-hsyn-Cre was purchased from addgene (#86641), and the human synapsin promoter was replaced with the CamKII promoter to generate **Lenti-CamKII-Cre**.

#### Viruses

The following viruses were purchased from addgene : AAV8-hsyn-hM3D(Gq)-mcherry #50474, AAVrg-EF1a-Cre #55636, AAVrg-Ef1a-DIO-Flp-WPRE-hGH #87306, AAV8.hSyn.DIO.mCherry #50459, AAV8-Ef1a-fDIO-mCherry #114471, AAV8.hsyn.DIO.eGFP.WPRE.bGH #50457, and AAV8.hSyn.hM3D.mCherry #50474.

### Quantification of activity-dependent transsynaptic

For experiments to quantify transsynaptic labeling with and without BACE, mCherry-expressing cells were counted within an identical 1 mm^2^ region over four consecutive 25 μm stained sections of Str. In all cases, the sections examined were roughly the same distance from the site of AAV injection. For experiments to quantify activity-dependent transsynaptic labeling, total GFP fluorescence minus background was measured identically in Str in sections that were equivalent distances from the injection site using ImageJ.

