## Supplementary material for "ATLAS: A rationally designed anterograde transsynaptic tracer": Figure s1

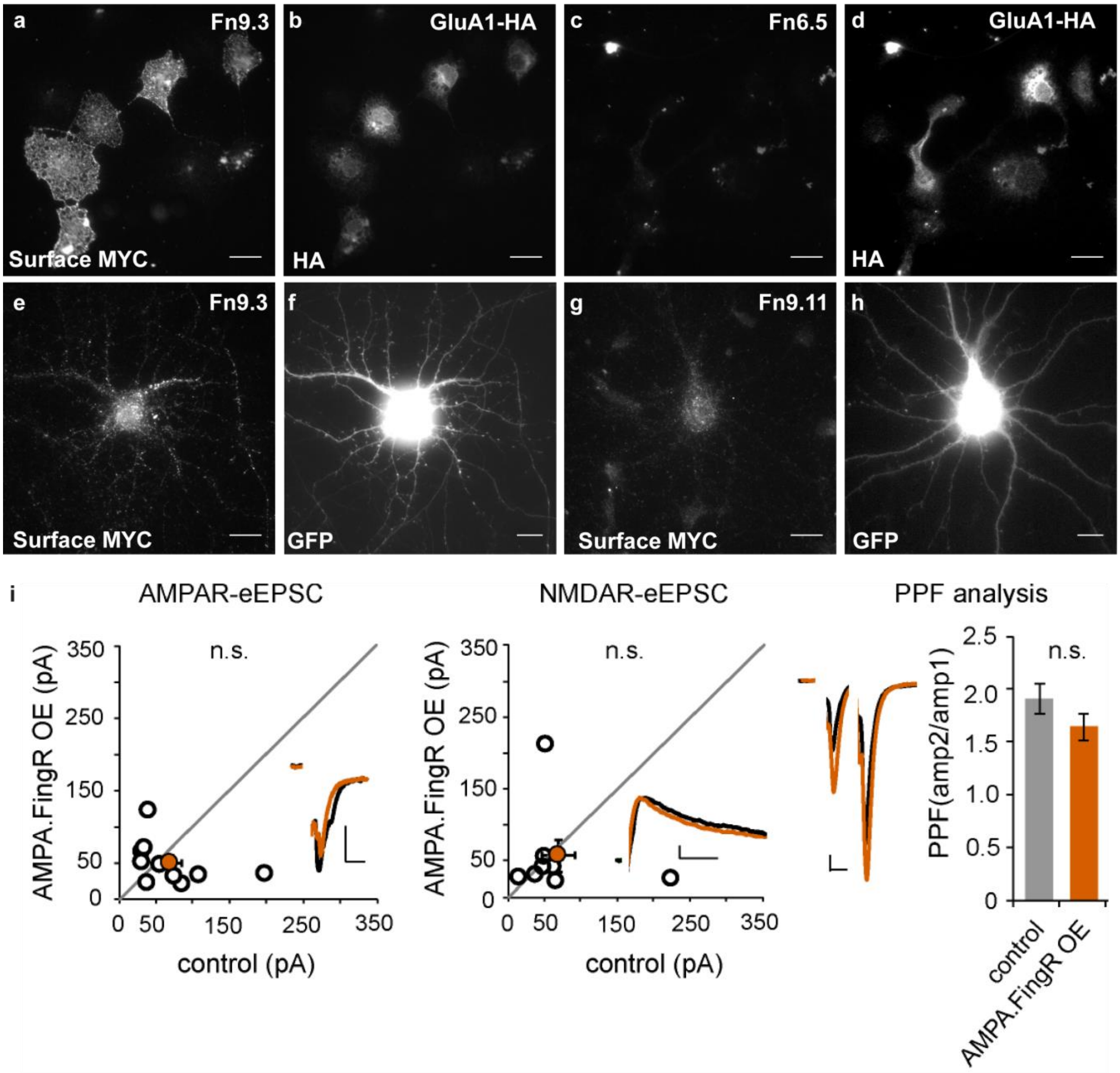


**Fig. s1 a-d** Screening of Fn clones in Cos7 cells. **a**., **b**. Surface labeling of MYC indicates binding of Fn9.3 to GluA1 in COS7 cells. **c**., **d**. Absence of surface MYC labeling indicates lack of binding of Fn6.5 to GluA1 in COS7 cells. **e**., **f**. MYC surface labeling following co-expression of ss-Fn9.3-MYC with GFP in cultured cortical neurons. **g**. **h**. Lack of MYC surface labeling following co-expression of ss-Fn9.1-MYC with GFP in cultured cortical neurons. Scale bar 20 μm **i**. Patch clamp recordings of CA1 neurons in hippocampal slices transfected biolistically with ss-AF-YFP indicate that there is no effect of AF expression on the amplitude of evoked AMPA- or NMDA receptor-mediated EPSCs, or on paired pulse facilitation compared to untransfected controls. Scale bars 20 pA and 20 ms for AMPA; 25 pA and 50 ms for NMDA; 20 pA and 20 ms for PPF.
